# The fate of carbon in a mature forest under carbon dioxide enrichment

**DOI:** 10.1101/696898

**Authors:** M. Jiang, B.E. Medlyn, J.E. Drake, R.A. Duursma, I.C. Anderson, C.V.M. Barton, M.M. Boer, Y. Carrillo, L. Castañeda-Gómez, L. Collins, K.Y. Crous, M.G. De Kauwe, K.M. Emmerson, S.L. Facey, A.N. Gherlenda, T.E. Gimeno, S. Hasegawa, S.N. Johnson, C.A. Macdonald, K. Mahmud, B.D. Moore, L. Nazaries, U.N. Nielsen, N.J. Noh, R. Ochoa-Hueso, V.S. Pathare, E. Pendall, J. Pineiro, J.R. Powell, S.A. Power, P.B. Reich, A.A. Renchon, M. Riegler, P. Rymer, R.L. Salomón, B.K. Singh, B. Smith, M.G. Tjoelker, J.K.M. Walker, A. Wujeska-Klause, J. Yang, S. Zaehle, D.S. Ellsworth

## Abstract

Atmospheric carbon dioxide enrichment (eCO_2_) can enhance plant carbon uptake and growth^1,2,3,4,5^, thereby providing an important negative feedback to climate change by slowing the rate of increase of the atmospheric CO_2_ concentration^6^. While evidence gathered from young aggrading forests has generally indicated a strong CO_2_ fertilization effect on biomass growth^3,4,5^, it is unclear whether mature forests respond to eCO_2_ in a similar way. In mature trees and forest stands^7,8,9,10^, photosynthetic uptake has been found to increase under eCO_2_ without any apparent accompanying growth response, leaving an open question about the fate of additional carbon fixed under eCO_2_^4, 5, 7,8,9,10,11^. Here, using data from the first ecosystem-scale Free-Air CO_2_ Enrichment (FACE) experiment in a mature forest, we constructed a comprehensive ecosystem carbon budget to track the fate of carbon as the forest responds to four years of eCO_2_ exposure. We show that, although the eCO_2_ treatment of ambient +150 ppm (+38%) induced a 12% (+247 gCm^-2^yr^-1^) increase in carbon uptake through gross primary production, this additional carbon uptake did not lead to increased carbon sequestration at the ecosystem level. Instead, the majority of the extra carbon was emitted back into the atmosphere via several respiratory fluxes, with increased soil respiration alone contributing ∼50% of the total uptake surplus. Our results call into question the predominant thinking that the capacity of forests to act as carbon sinks will be generally enhanced under eCO_2_, and challenge the efficacy of climate mitigation strategies that rely on CO_2_ fertilization as a driver of increased carbon sinks in standing forests and afforestation projects.

## Main text

Globally, forests act as a large carbon sink, absorbing ∼30% of total anthropogenic CO_2_ emissions^1, 12^, an ecosystem service that has tremendous social and economic value. Whether mature forests will remain carbon sinks into the future is of critical importance for aspirations to limit climate warming to no more than 1.5 °C above pre-industrial levels^13^. Free-Air CO_2_ Enrichment (FACE) experiments provide an opportunity to determine the capacity of ecosystems to sequester carbon under the higher atmospheric CO_2_ concentrations expected in the future^3,4,5, 7, 8, 10, 11^. Evidence gathered from the four first generation forest FACE experiments, which all measured responses of rapidly-growing young forest plantations, has generally indicated a strong CO_2_ fertilization effect on biomass growth^3, 4^. This CO_2_ fertilization effect has been hypothesized to be one of the largest drivers of the terrestrial carbon sink and its acceleration in recent decades^14^, potentially accounting for up to 60% of present-day terrestrial carbon sequestration^2^. Given that younger trees are generally more responsive to rising CO_2_ than mature trees^11^, extrapolating evidence collected from these experiments may be argued to provide an upper limit on how much carbon can be stored by global forests under eCO_2_^15^. However, evidence from experiments with older trees suggests that although eCO_2_ increases leaf photosynthesis to a similar degree as in young forests, stimulation of biomass growth and carbon storage may be lower or absent^7,8,9,10^. Reconciling these conflicting observations is a crucial step towards quantifying the carbon sequestration capacity of mature forests in the future. It requires that we identify the fate of the extra carbon fixed under eCO_2_ in these complex ecosystems, which are expected to be closer to a state of equilibrium between carbon uptake and turnover, compared to young growing stands.

The *Eucalyptus* FACE (EucFACE) experiment is the world’s first replicated, ecosystem-scale mature forest FACE experiment (Extended Data Figure 1). It is established in a warm-temperate evergreen forest that has remained undisturbed for the past 90 years and that is dominated by regionally widespread tree *Eucalyptus tereticornis*. The site is characterized by soils of low fertility with an understorey dominated by native grasses and shrubs. Seven ecosystem-scale models were used to predict the eCO_2_ response at EucFACE in advance of the experiment^16^, highlighting three alternative hypotheses for the expected ecosystem response based on plausible assumptions incorporated in different models^17^. These hypotheses were: (i) enhanced photosynthesis under eCO_2_ would lead to increased biomass accumulation; (ii) eCO_2_-induced increase in photosynthesis would be directly down-regulated by limited nutrient availability; or (iii) eCO_2_-induced increase in photosynthesis would lead to increased autotrophic respiration^16^. This range of predictions among a suite of well-tested models indicated a prognostic knowledge gap as to how the carbon cycling of mature forests would respond to the expected rise in CO_2_ concentration^11^, which is crucial to resolve in the face of future carbon-climate uncertainty^18^.

**Figure 1.**
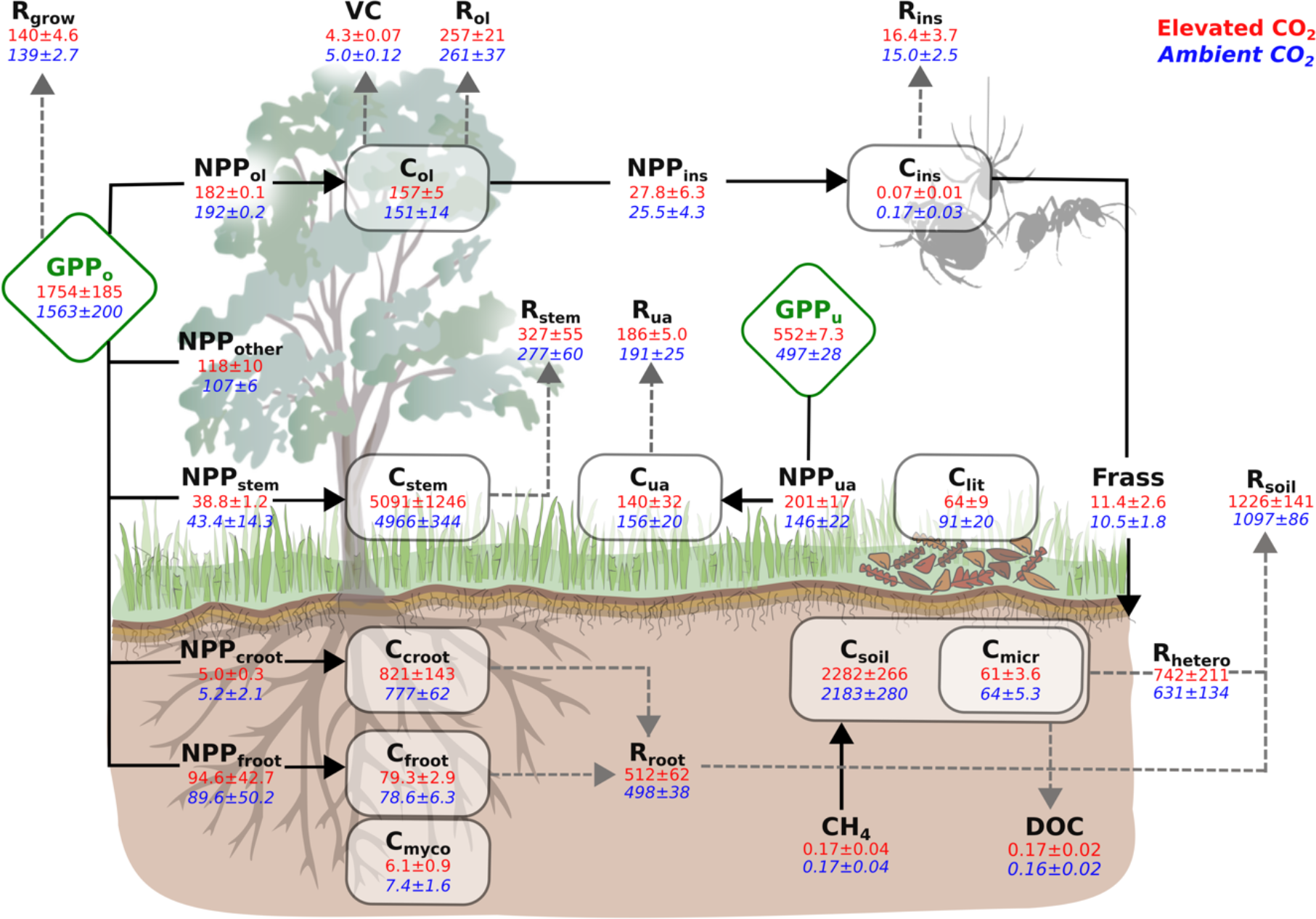
A comprehensive carbon budget under ambient and elevated CO_2_ treatment in a mature forest ecosystem. Diamond boxes are gross primary production for overstorey (GPP_o_) and understorey (GPP_u_), respectively. Squared boxes are carbon stocks (gCm^-2^), including overstorey leaf (C_ol_), stem (C_stem_), coarse root (C_croot_), fineroot (C_froot_), understorey aboveground (C_ua_), leaf litter (C_lit_), soil (C_soil_), microbe (C_micr_), aboveground insect (C_ins_), and mycorrhizae (C_myco_). Unboxed variables are carbon fluxes (gCm^-2^yr^-1^), including net primary production of overstorey leaf (NPP_ol_), stem (NPP_stem_), coarse root (NPP_croot_), fineroot (NPP_froot_), and understorey aboveground (NPP_ua_), overstorey leaf consumption by insects (NPP_ins_), respiration fluxes of overstorey leaf (R_ol_), stem (R_stem_), root (R_root_), understorey aboveground (R_ua_), growth (R_grow_), insect (R_ins_), heterotroph (R_hetero_), and soil (R_soil_), and volatile carbon emission (VC), frass production (Frass), dissolved organic carbon (DOC), and soil methane net uptake (CH_4_). Solid arrow lines are fluxes entering a pool, dotted arrow lines are fluxes leaving a pool. Blue italic values are means ± one standard deviation of the ambient CO_2_ treatment (n=3), whereas red values are means ± one standard deviation of the elevated CO_2_ treatment (n=3). All values are normalized by a linear mixed-model with plot-specific pre-treatment leaf area index as a covariate to account for pre-existing differences. Summary of variable definitions and data availability is provided in Extended Data Table 1.

To date, both canopy trees and understorey plants at EucFACE have shown increased rates of leaf photosynthesis but the canopy trees showed no significant increase in aboveground biomass growth under eCO_2_^7^, reflecting a similar lack of response observed in other eCO_2_ experiments on mature trees^8,9,10^. Incorporating leaf-scale gas exchange measurements into a process-based tree stand model, it was estimated that the observed +19% stimulation of light-saturated overstorey leaf photosynthesis^7^ corresponded to a +12% stimulation of whole-canopy gross primary production (GPP) response to eCO_2_^19^. However, the probable fate of the extra carbon fixed under eCO_2_ remained undetermined. Where did the extra carbon go?

To answer this question, we compiled measurements on all major carbon pools and fluxes collected over four years of experimental treatment (2013-2016), including individual and aggregated biomass and associated fluxes measured or inferred from plants, litter, soil, microbes, and insects, and constructed an ecosystem carbon budget (Figure 1) under both ambient (aCO_2_) and eCO_2_ conditions (+150 ppm). We first confirmed mass balance of the ecosystem carbon budget by checking agreement between independent estimates of GPP and soil respiration (R_soil_) derived from separate data streams (Extended Data Figure 2; see Methods). For GPP of the aCO_2_ plots, we confirmed that a process-based model estimate of overstorey and understorey GPP (2059 ± 211 gCm^-2^yr^-1^), driven by site-specific meteorology and physiological data, agreed with the sum of data-driven estimates of net primary production (NPP) and autotrophic respiration (1968 ± 80 gCm^-2^yr^-1^). The carbon-use efficiency (NPP/GPP) of this mature forest was estimated to be 0.29 ± 0.02, which is on the low end of global forest estimates, but consistent with studies that have found this ratio tends to decline with stand age^20^. We further confirmed carbon mass balance for R_soil_ of the aCO_2_ plots by comparing soil chamber-based estimates (1097 ± 86 gCm^-2^yr^-1^) with the sum of litterfall and independently estimated root respiration (1036 ± 27 gCm^-2^yr^-1^), assuming no change in soil carbon pool (see Methods). This agreement between independent estimates of components of the ecosystem carbon budget gives confidence that our measurements captured the pools and fluxes of carbon with low aggregate uncertainty and hence allows us to infer the fate of the extra carbon fixed under eCO_2_.

**Figure 2.**
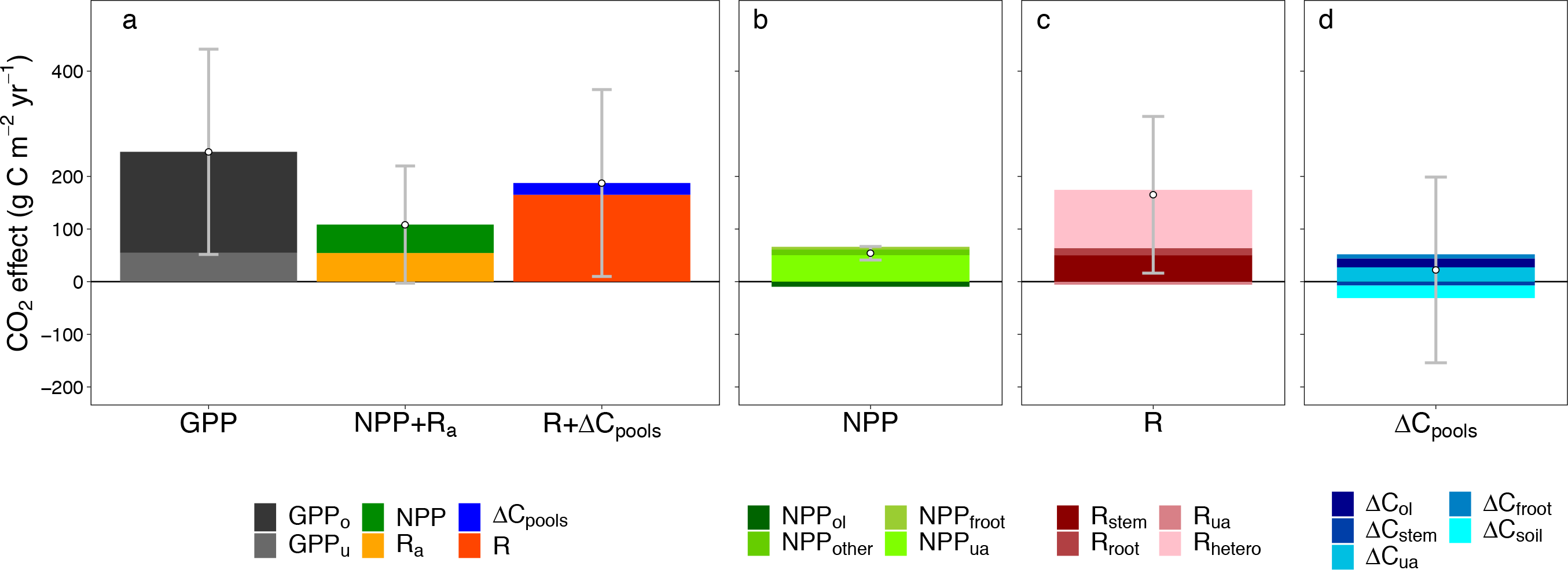
The fate of additional carbon fixed under elevated CO_2_ (eCO_2_) in a mature forest ecosystem. **a)** Column **“**GPP” represents the total eCO_2_-induced increases in overstorey and understorey gross primary production (GPP_o_ and GPP_u_, respectively), “NPP + R_a_” represents the sum of net primary production and autotrophic respiration response, “R + ΔC_pools_” represents the sum of ecosystem respiration and carbon storage response. **b)** The relative contributions of individual NPP fluxes to the aggregated NPP response to eCO_2_, including NPP responses of overstorey leaf (NPP_ol_), twigs, barks and seeds (NPP_other_), fineroot (NPP_froot_), and understorey aboveground (NPP_ua_); **c)** The relative contributions of individual respiratory fluxes to the aggregated R response to eCO_2_, including respiration responses of stem (R_stem_), root (R_root_), understorey aboveground (R_ua_), and soil heterotroph (R_hetero_); and **d)** The relative contributions of individual change in carbon storage to the aggregated ΔC_pools_ response to eCO_2_, including changes in pool of overstorey leaf (ΔC_ol_), stem (ΔC_stem_), understorey aboveground (ΔC_ua_), fineroot (ΔC_froot_), and soil (ΔC_soil_). Variables with an absolute mean CO_2_ effect of < 5 gCm^-2^yr^-1^ are excluded from the figure for better visual clarification. Individual CO_2_ responses are reported in Extended Data Figure 4. Each color represents the CO_2_ response of a flux variable, point indicates the net sum of all variables for a column, and the grey error bar represents one standard deviation of the estimated column sum at the plot-level (see Methods). The CO_2_ effect is estimated using a linear mixed-model analysis with plot-specific pre-treatment leaf area index as a covariate to account for pre-existing differences (see Methods). The un-normalized response is provided in Extended Data Figure 3, which generally agrees with findings present in this figure, but with less statistical precision.

To accommodate the inherent pre-treatment plot differences (see Methods), we normalized the CO_2_ responses across plots by using a linear mixed-model with plot-specific pre-treatment leaf area index as a covariate^21, 22^. The un-normalized eCO_2_ responses are provided in Extended Data Figure 3, and generally confirm the findings but with less statistical precision. Our normalized responses (Figure 2, Extended Data Figure 4) showed that eCO_2_ induced an average of 12% increase (+247 ± 195 gCm^-2^yr^-1^, mean ± one standard deviation) in carbon uptake, including contributions of overstorey (+192 ± 157 gCm^-2^yr^-1^) and understorey GPP (+55 ± 17 gCm^-2^yr^-1^). The fate of this additional carbon entering the system under eCO_2_ was primarily traced to an increase in R_soil_ (+128.8 ± 95.2 gCm^-2^yr^-1^, or 52% of the carbon uptake surplus), followed by a smaller increase in stem respiration (R_stem_; +50.2 ± 47.2 gCm^-2^yr^-1^, or 20% of the carbon uptake surplus). In comparison, the increase in total NPP (+54 ± 12.9 gCm^-2^yr^-1^, or 22% of the carbon uptake surplus) was similar in magnitude to the increase in R_stem_, but the increase in storage of the total carbon pools at the ecosystem-level was much smaller (ΔC_pools_; +22.3 ± 176.4 gCm^-2^yr^-1^, or 9% of the carbon uptake surplus). There was thus little evidence of additional carbon accumulation under eCO_2_ in this mature forest ecosystem. We then compared three alternative methods (see Methods) of estimating net ecosystem production (NEP; Figure 3). All three indicated that the ecosystem remained close to carbon-neutral under ambient CO_2_ over the experimental period (mean ± SD for the methods: 74 ± 258, -35 ± 142, 115 ± 96 gCm^-2^yr^-1^, respectively), and that eCO_2_ of +150 ppm did not result in statistically significant increases in ecosystem carbon storage (149 ± 261, -92 ± 216, 137 ± 230 gCm^-2^yr^-1^, respectively).

**Figure 3.**
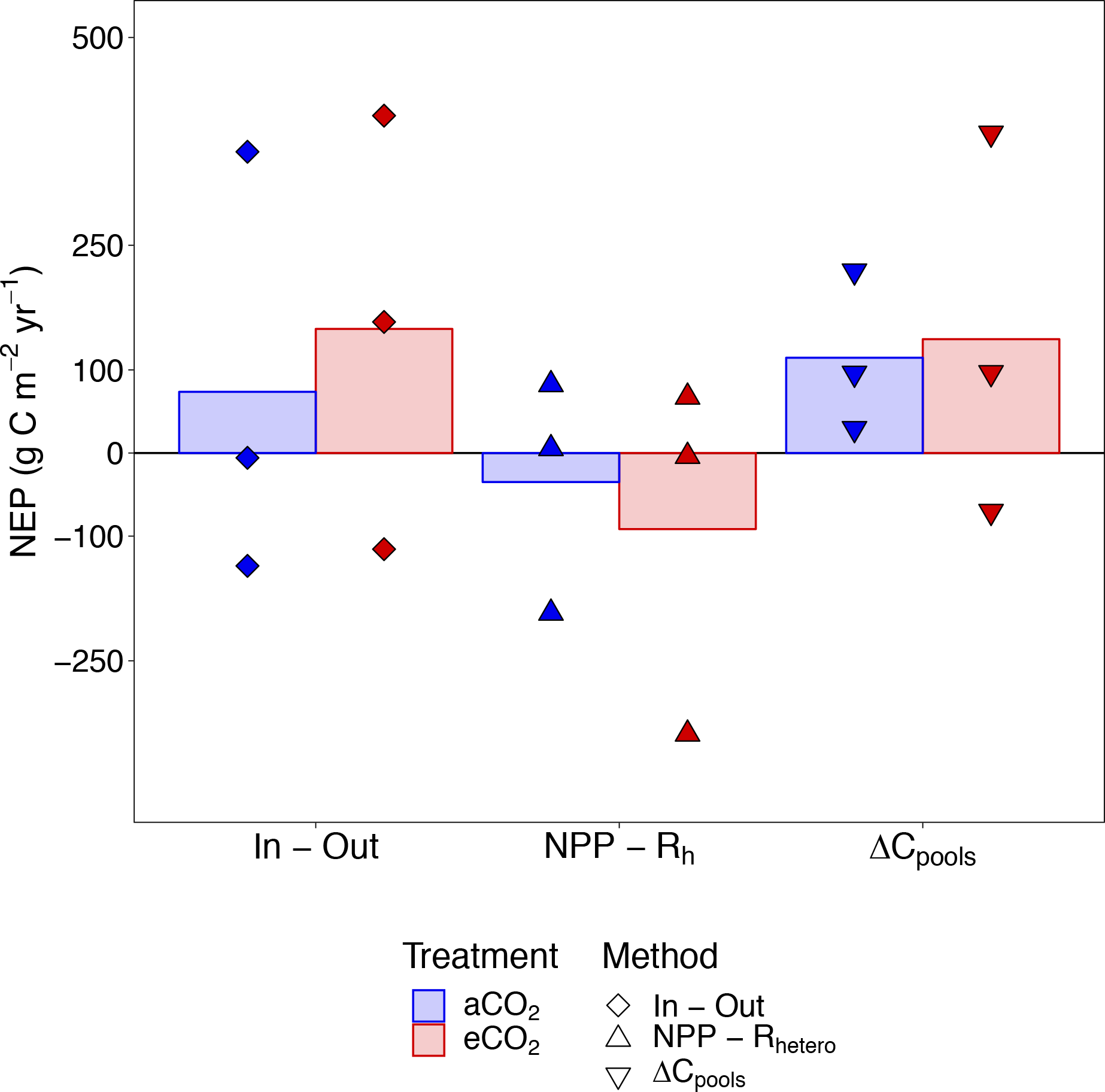
Estimates of net ecosystem production (NEP) under ambient and elevated CO_2_ treatment at EucFACE. Positive values indicate ecosystem net carbon uptake by the ecosystem. “In - Out” calculates NEP based on the difference between total influxes and total outfluxes. “NPP - R_hetero_” calculates NEP based on the difference between net primary production (NPP) and heterotrophic respiration (R_hetero_). “ΔC_pools_” derives NEP based on incremental changes in all ecosystem carbon pools. Colored bars indicate treatment means based on each method (n=3), with blue representing ambient and red representing elevated CO_2_ treatment. Individual dots are plot-level NEP, derived based on different methods (see Methods). Values are normalized by a linear mixed-model with plot-specific pre-treatment leaf area index as a covariate to account for pre-existing differences. Horizontal dotted line indicates NEP equals zero.

The relatively small but positive NPP response to eCO_2_ was mainly driven by the understorey aboveground NPP response (NPP_ua_; +50.3 ± 14.6 gCm^-2^yr^-1^), which was 93% of the net NPP response (Figure 2). However, this significant NPP_ua_ response did not result in an equivalent eCO_2_ effect on understorey aboveground biomass increment (+27.2 ± 24.2 gCm^-2^yr^-1^), suggesting a possible higher understorey biomass turnover under eCO_2_. Smaller fluxes, often neglected in other ecosystem carbon budgets, such as leaf consumption by insect herbivores (NPP_ins_; 25.5 ± 4.3 vs. 27.8 ± 6.3 gCm^-2^yr^-1^, aCO_2_ vs. eCO_2_ mean ± SD), insect frass production (Frass; 10.5 ± 1.8 vs. 11.4 ± 2.6 gCm^-2^yr^-1^), vegetation volatile carbon emission (VC; 5.0 ± 0.12 vs. 4.3 ± 0.07 gCm^-2^yr^-1^), net ecosystem methane uptake (CH_4_; 0.17 ± 0.04 vs. 0.17 ± 0.04 gCm^-2^yr^-1^), and leaching of dissolved organic carbon (DOC; 0.16 ± 0.02 vs. 0.17 ± 0.02 gCm^-2^yr^-1^), contributed to the closure of the overall ecosystem carbon budget (Figure 1; Extended Data Figure 2), but were not important in explaining pathways of the carbon uptake surplus under eCO_2_ (Figure 2, Extended Data Figure 4).

Here we provide some of the first replicated experimental evidence on the probable fate of carbon under eCO_2_ in intact mature forests. We found that increased R_soil_ accounted for ∼50% of the extra photosynthate produced by plants under eCO_2_. It has been suggested that the increase in R_soil_ at EucFACE was likely a consequence of increased root and rhizosphere respiration^23, 24^, in contrast to other FACE sites where increased R_soil_ was attributed to enhanced soil organic matter decomposition (e.g. DukeFACE^25^). Here, the eCO_2_-induced increase in R_soil_ was not accompanied by substantial changes in pools of fine root (+7.9 ± 8.4 gCm^-2^yr^-1^), microbial (+2.5 ± 2.9 gCm^-2^yr^-1^), mycorrhizal (+0.5 ± 0.4 gCm^-2^yr^-1^), leaf litter (-1.7 ± 6.2 gCm^-2^yr^-1^) or soil carbon (-23.8 ± 130.3 gCm^-2^yr^-1^), suggesting that the additional carbon fixed under eCO_2_ may have led to an enhanced carbon transport belowground and a rapid belowground turnover of this flux. An initial enhancement in nitrogen and phosphorus mineralization was observed^26^, which suggested that the increased R_soil_ with eCO_2_ could reflect soil organic matter priming with the potential to alleviate plant nutrient stress in this phosphorus-deprived environment^26, 27^. However, the enhanced soil mineralization rate and associated increase in nutrient availability did not persist over time^26^, indicating that this increased belowground carbon allocation and the rapid turnover of this flux was not effective in increasing phosphorus availability to the plants^7,28^.

The ecosystem carbon budget presented here provides an opportunity to confront the three alternative hypotheses of the response of this system to eCO_2_ treatment that emerged from model predictions made in advance of the experiment^16^. Our data do not support any of the three hypotheses. The eCO_2_-induced increase in photosynthesis was not strongly down-regulated by low nutrient availability; nor did the eCO_2_-induced additional carbon uptake lead to additional biomass accumulation, or enhanced aboveground respiration^16^. These predictions reflect common mechanisms by which terrestrial vegetation models implement nutrient limitation of the eCO_2_ response^16, 17, 29, 30^. In contrast, our results suggest a direct connection between plant photosynthesis and belowground activity, in which increased belowground carbon allocation increased soil respiration at a rate that accounted for half of the extra carbon fixed under eCO_2_. This increased soil respiration has been demonstrated by some models to be an important and often overlooked mechanism that reduces global soil carbon sequestration relative to estimates by many current models^31^. As a consequence of including this rapid turnover of the increased belowground carbon allocation in terrestrial biosphere models, the time lag in emitting some of the extra carbon via biomass accumulation and litterfall input into the soils may be reduced, thereby leading to faster cycling of carbon^32^ and therefore possible different trajectories of carbon-climate predictions for the future.

A major form of land-based climate mitigation actions envisaged in the Paris Agreement is to enhance forest biomass carbon stocks globally through the protection of existing, largely mature, forests, and through afforestation of new areas. The mitigation potential of forests lies in the accumulated stock of ecosystem carbon, not in the short-term rate of forest photosynthesis. The probable fate of additional carbon determined in our study (Figure 2) challenges the current thinking that non-aggrading mature forests can contribute to enhanced carbon sinks due to CO_2_ fertilization^33^, which further questions the allowable CO_2_ emission targets sourced from existing carbon cycle models^13, 34^. Given that the effect of CO_2_ fertilization may be one of diminishing returns over time^14^, the statistically insignificant eCO_2_ effect on NEP (Figure 3), if representative of mature forest ecosystems generally, suggests an even weaker carbon sink in the future, especially in low fertility systems such as EucFACE. Future research efforts should target a deeper understanding of the nutrient-carbon feedbacks that likely constrain the carbon sink potential of mature forests under eCO_2_, and evaluate the implications of a potentially weaker terrestrial land carbon sink in the development of robust mitigation strategies in the face of climate change.

## Methods

### EucFACE site description

The EucFACE facility (Extended Data Figure 1) is located in a mature evergreen *Eucalyptus* forest on an alluvial spodosol in western Sydney, Australia (33°36’S, 150°44’E). The site has been a remnant patch of native Cumberland Plain woodland since the 1880’s and has remained unmanaged for at least the past 90 years, with *Eucalyptus tereticornis* Sm. as the dominant tree species. *Eucalyptus* trees occur naturally across Australia, accounting for 78% of native forest area in Australia^35^ and are planted widely around the globe^36^. Infrastructure for six large circular plots (490 m^2^ each) was established in 2010. Starting on 18^th^ September 2012, three plots were subjected to free-air CO_2_ enrichment treatment using computer-controlled pre-dilution method. The CO_2_ concentrations at EucFACE were ramped up over a six-month period, increasing by +30 ppm every five weeks in discrete steps (+30, 60, 90, 120, and 150 ppm). The full elevated CO_2_ treatment of +150 ppm started on 6^th^ February 2013 during daylight hours over all days of the year. The site is characterized by a humid temperate-subtropical transitional climate with a mean annual temperature of 17.5°C and a mean annual precipitation of 800 mm (Figure S1). The soil is a Holocene alluvial soil of low-fertility with low phosphorus content^7,37^. Soil texture is a loamy sand (> 75% sand content) up to 50 cm in depth. From ca. 50 to 300 cm depth, soils are sandy clay loam, with > 30% silt and clay. Average bulk density is 1.39, 1.69 and 1.71 g cm^-3^ for depths of 0-10, 10-20 and 20-30 cm, respectively (Figure S2). Permanent groundwater depth is ∼11 m below the soil surface^38^. Understorey vegetation is a diverse mixture of 86 species including forbs, graminoids and shrubs^39^. The dominant understorey species is *Microlaena stipoides*, a C3 perennial grass that accounted for ∼70 % of herbaceous biomass, and responded rapidly to rainfall variability^40^.

### Estimates of carbon pools and fluxes

We estimated plot-specific carbon pools and fluxes at EucFACE over 2013-2016 (Extended Data Table 1). We defined pools as a carbon reservoir and annual increments as the annual change in the size of each reservoir. We compartmentalized the ecosystem into 10 carbon pools, namely overstorey leaf (C_ol_), stem (C_stem_), fine root (C_froot_), coarse root (C_croot_), understorey aboveground (C_ua_), soil (C_soil_), microbe (C_micr_), mycorrhizae (C_myco_), leaf litter (C_lit_), and aboveground insect (C_ins_) carbon pools, and reported pool size in the unit of gCm^-2^. We defined fluxes as components of the carbon flow through the system, and report them in the unit of gCm^-2^yr^-1^. All annual incremental changes in carbon pools were reported in gCm^-2^yr^-1^ with a symbol Δ. We converted estimates of biomass into carbon content using variable-specific carbon fractions (f) defined in Extended Data Table 2. Below we describe how each pool and flux was estimated.

#### Pools

**Soil carbon pool (C_soil;_** Figure S2) was estimated based on quarterly sampled soil carbon content (oven-dried at 40 °C for 48 hours) and plot-specific soil bulk density at three depths (0 - 10 cm, 10 - 20 cm, 20 - 30 cm). Out of the 15 dates when samples were taken, only 3 of these measured soil carbon content below the top 10 cm of soil. To obtain a more accurate estimate of annual incremental change in soil carbon pool, we therefore reported soil carbon pool for the top 10 cm only. There were no temporal and eCO_2_ trends in soil carbon content at deeper depths.

**Overstorey leaf carbon pool (C_ol_;** Figure S3) was estimated based on continuous measures of leaf area index (LAI) and specific leaf area (SLA), following C_ol_ = LAI × SLA × f_ol_, where f_ol_ is a carbon fraction constant for overstorey leaf (Extended Data Table 2). Daily averages of plot-specific LAI were estimated based on the attenuation of diffuse radiation in a homogenous canopy^22^. The number of observations varies between days, depending on the number of 30-minute cloudy periods. SLA was estimated based on time-series measures of leaf mass per area (LMA), and was then linearly interpolated to plot-specific daily values over time.

**Stem carbon pool (C_stem_;** Figure S4) was estimated based on tree-specific height and diameter at breast height (DBH) measurements, and an allometric scaling relationship derived based on *E. tereticornis*^7,41^. DBH changes were measured repeatedly at roughly one month intervals at 1.3 m height. Bark was periodically removed from under the dendrometer bands - this effect on DBH was considered by calculating biomass once per year using December data only. Stem biomass data were summed for each plot and averaged over the plot area to obtain ground-based estimates, and was then converted into C_stem_ using treatment-specific carbon fraction (Extended Data Table 2).

**Understorey aboveground carbon pool (C_ua_**; Figure S5) was estimated at 1-3 month intervals between February 2015 and December 2016 using non-destructive measurements of plant height obtained from stereo-photography^42^. In each of the four 2m × 2m understorey monitoring subplots within each plot, stereo photographs were collected using a Bumblebee XB3 stereo camera (Point Grey Research) mounted ∼2.4 m above the ground surface and facing vertically downwards towards the center of the subplot. Stereo images were taken at dusk under diffuse light conditions to avoid measurement errors related to shadows from trees and EucFACE infrastructure. On each sampling date, three sets of stereo photographs were taken in each subplot to produce large number (i.e. 100,000s) of understorey plant height estimates from which mean plant height (H_mean_, in m) was calculated for each plot. Understorey aboveground biomass (B_ua_, in kg m^-2^) for each plot was predicted from H_mean_ using an empirical model developed for the grassy understorey vegetation at EucFACE (B_ua_ = 1.72 * H_mean_ – 0.05)^42^. The four subplot-level estimates were averaged to obtain a plot-level estimate of B_ua_, and then converted to an estimate of C_ua_ using a carbon fraction constant (Extended Data Table 2).

**Root carbon pool (C_root_)** consists of fineroot (C_froot_) and coarseroot (C_croot_) pools, with C_froot_ defined as roots with diameter < 2 mm, with the remaining roots or woody roots defined as C_croot_ (Figure S6). The C_root_ pool includes roots of both overstorey and understorey vegetation. Total root carbon pool (C_root_) was estimated based on an allometric relationship between root biomass (B_root_) and stand basal area (derived from DBH) derived for Australian forest species^43^, as follows:

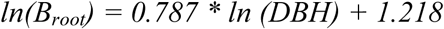

Fineroot biomass was estimated based on standing biomass sampled at 4 subplots per plot at 2 depths (0 - 10 cm and 10 - 30 cm) over the period of 2014-2015^27^. Plot-specific fineroot biomass was taken by summing biomass data across depths. Coarseroot biomass was estimated as the net difference between fineroot and total root biomass. The fineroot and coarseroot biomass were multiplied by the corresponding carbon fraction constants to obtain C_froot_ and C_croot_, respectively (Extended Data Table 2).

**Microbial carbon pool (C_micr_)** was estimated based on fumigation extraction and 0.5 M K_2_SO_4_ extraction as in Ref. 23 using samples taken at 0-10 cm soil depth over the period of 2012 - 2015. Total organic carbon was determined on a Shimadzu TOC analyzer (TOC-L TNM-L; Shimadzu, Sydney, Australia), which was then multiplied by soil bulk density over the same soil depth to obtain the C_micr_ (Figure S7a).

**Mycorrhizal carbon pool (C_myco_)** for the top 10 cm of soil was estimated via measurements of colonization of mycorrhizal in-growth bags, carbon isotopic partitioning, microbial phospholipid fatty acid abundance and C_micr_. Nine 45 µm nylon mesh bags (4 × 5 cm) filled with sand, which excluded roots but allowed access of fungi^44^, were buried in November 2014 in each experimental plot and three bags were subsequently collected every four months for one year. Phospholipid-derived fatty acids (PLFA), a proxy for total microbial biomass abundance, were quantified in sand bags and native field soil following the protocol by Ref. 45 δ^13^C values of ground subsamples of this sand, native soil carbon, and aboveground plant tissue (leaves of Eucalypts in April 2014) were used to estimate the fraction of the accumulated carbon in sand bags that was derived from plant carbon using isotopic mass balance. Due to the exclusion of roots, plant derived carbon in bags can be attributed to mycorrhiza. This plant-derived unitless fraction was then multiplied by the total concentration of PLFA in sand bags to obtain the amount of the total PLFA contributed by mycorrhiza (µg PLFA / g sand). To scale this to native soil PLFA concentrations we then calculated the ratio between mycorrhizal PLFA in sand bags to total PLFA in soil (representing the total microbial pool). Subsequently, to estimate C_myco_, this ratio was multiplied by the C_micr_ in each plot (Figure S7b).

**Leaflitter carbon pool (C_lit_)** was estimated based on leaf litter decomposition rate and leaf litterfall data collected by litter baskets (Figure S8)^22^. Leaf litter decomposition rates were estimated over 24 months using litter bags. Briefly, 2 g air-dried *Eucalyptus* litter was added to 10 × 15 cm litter bags with a 2-mm mesh size. Twelve litter bags were randomly allocated to 4 subplots within each treatment plot, and two litter bags were collected at 3, 6, 9, 12, 18 and 24 months to calculate mass loss over time (mass loss was averaged across the two replicates from each subplot). A leaflitter exponential decay function was estimated for each plot, based on data collected over this 24-month period. Leaf litterfall was estimated from monthly collections of material from circular fine-mesh traps (each 0.2 m^2^) at 8 random locations for each plot. We then applied the exponential decay function with litterfall biomass to obtain C_lit_, assuming a carbon fraction constant (Extended Data Table 2).

**Insect carbon pool (C_ins_)** was estimated based on two different sampling techniques, with aerial insects partially estimated based on monthly dead insect data collected from circular fine-mesh traps of 0.2 m^2^ at 8 random locations for each plot^46^, and understory insects estimated based on vacuum suction sampling from 2 locations for each plot^47^. The vacuum suction method collected invertebrates from understorey vegetation in two 1 x 1 m subplots using a petrol-powered ‘G-Vac’ vacuum device run on full-throttle for 20 s, for a total of 5 sampling campaigns. Trapping locations were randomly chosen and fixed between sampling campaigns. All invertebrates were sorted from debris, dried to constant weight at 60°C and weighed on a microbalance with an accuracy of 1 µg. We assume that vacuum samples as well as litter trap samples represent point estimates of invertebrate abundance. Then, the total biomass of sampled invertebrates was summed across sampling methods within each plot. A constant carbon fraction based on Ref. 48 (Extended Data Table 2) was used to convert biomass into C_ins_ pool (Figure S9).

#### Ecosystem carbon uptake fluxes

**Overstorey gross primary production (GPP_O_)** for each plot was provided by a stand-level model simulation (MAESTRA), forced by hourly meteorological data and interpolated photosynthetic parameters measured at the site (Figure S10a)^19^. In MAESTRA, each plot consists of individual tree crowns that are located and parameterized with measured coordinates, crown size, and LAI. Each crown was divided into six layers, with leaf area uniformly distributed into each layer. Within each layer, the model simulated twelve points.

The radiation at each grid point considered shading from upper crown and surrounding trees, solar angle (zenith and azimuth), and light source (diffused or direct). According to the radiation, the leaf area at each grid point was divided into sunlit and shaded leaves, which was used to calculate gas exchange using a Farquhar^49^ type formulation for photosynthesis. Calculations for carbon flux were parameterized with *in situ* leaf gas exchange measurements^7,50^. Respiration and its temperature dependence were also quantified using data collected on site. The output was evaluated against measured canopy-scale transpiration data^19^.

Similarly, **understorey GPP (GPP_u_)** (Figure S10b) was simulated using MAESTRA with photosynthetic parameters taken for the grass *Microlaena stipoides*^40^. The parameterization of understory vegetation is different from that of the canopy. In each plot, the understory was assumed to form a single crown covering the whole plot (i.e., a circle with 12.5 m radius) at a height of 1.5 m. The LAI of the understory was estimated using phenology camera digital photographs taken at four permanent understorey vegetation monitoring subplots in each plot^42^. The average green pixel content was calculated from three photos in each subplot, and assumed to be the same as the fraction of absorbed PAR. We then assumed a light extinction coefficient of 0.5 in Beers’ Law and calculated understorey LAI. Before 2014 there were 3 campaigns per year while from 2014 the cameras were automated, and we used the fortnightly averages. Leaf gas exchange parameters were obtained from Ref. 40 and covered four to six campaigns per year from 2013 to 2016. We estimated a one-time g_1_ parameter^51^ for all plots and time, and assumed constant carboxylation rate (V_cmax_) and electron transport rate (J_max_) values at 25 °C across plots. Basal leaf respiration rate and the temperature dependence of photosynthesis and respiration were assumed to be the same as the canopy. The understory simulation was conducted separately from the canopy, with canopy LAI from Ref. 22 included to account for the shading from the canopy, branches and stems on the understory.

For the **methane net flux (CH_4_),** air samples were collected following the closed-chamber method (or Non-Flow-Through Non-Steady-State [NFT-NSS] method). Seven replicated chambers were available for each plot. Headspace samples were collected monthly, over a period of one hour and analyzed by gas chromatography. Fluxes were estimated by a mixture of linear and quadratic regressions (depending on goodness-of-fit), assuming a constant air pressure of one atm and correcting the air temperature inside the chambers for each air sample^52^. The CH_4_ fluxes are net fluxes, which represent the sum of: 1) CH_4_ efflux (emissions from the soil into the atmosphere); 2) CH_4_ influx (uptake from the atmosphere into soil). Here, the annual net CH_4_ flux was an ecosystem influx and was presented as positive values (Figure S11a).

#### Production fluxes

Plant **net primary production (NPP)** is the sum of overstorey leaf (NPP_ol_), stem (NPP_stem_), fine root (NPP_froot_), coarse root (NPP_croot_), other (including twigs, barks, and seeds; NPP_other_), understorey aboveground (NPP_ua_), and consumption of overstorey leaf by insect herbivores (NPP_ins_). NPP_ol_ and NPP_other_ were estimated based on monthly litter data collected from circular fine-mesh traps of 0.2 m^2^ at eight random locations for each plot (Figure S12). Litter were sorted into leaf, twigs, bark, and seeds, dried to constant mass at 40 °C and weighed. A subsample was reweighed when dried to constant mass at 70 °C and a small moisture correction was applied to the leaf component of the whole dataset. NPP_ol_ was computed as the sum of annual leaf litter, which excluded leaf consumption by insects. For twigs, we assumed strictly annual turnover across the years. NPP_stem_ (Figure S13) and NPP_croot_ (Figure S14) were estimated based on annual incremental change of stem biomass and coarse root biomass, respectively. NPP_froot_ was estimated based on samples collected from the in-growth cores at 4 different locations per plot (Figure S14).

NPP_ua_ was estimated based on biomass clippings taken between 2015 - 2017, assuming one understorey turnover per harvest interval (Figure S15). We used a clip-strip method of biomass harvest as has been applied previously at the BioCON experiment^53^. Specifically, four narrow strips, each with a size of 1 m x 0.1 m, were situated in each of the experimental plots at least 2 m away from the vertical pipes for FACE, while avoiding the understory shrubs. The understory herbaceous species were clipped approximately 1 cm above soil level. The total mass per harvest represents the total production. Biomass samples were oven dried for two days at 60 °C, and converted into carbon mass by applying a constant fraction (Extended Data Table 2).

NPP lost to overstorey leaf consumption by insect herbivores (NPP_ins_) was estimated based on insect frass data (Frass) collected from the circular fine-mesh traps, and a relationship between frass mass and insect consumed leaf mass derived based on multiple *Eucalyptus* tree species at different CO_2_ concentrations (Figure S16a)^54, 55^. Frass was estimated based on annual collection of frass biomass collected from the circular fine-mesh litter traps with their associated carbon content (Extended Data Table 2; Figure S16c).

#### Outfluxes

Leaching lost as **dissolved organic carbon (DOC)** from soils was estimated based on concentrations of DOC in soil solutions, provided by water suction lysimeter measurements^26^. Lysimeters were installed to two depths (0 - 15 cm and 35 - 75 cm, which is immediately above the impermeable layer). Here we assumed that DOC reaching deeper depth is lost from the system at a rate of 20 ml m^-2^ d^-1^, which is an estimate of the daily drainage rate at the site (Figure S11b).

**Plant autotrophic respiration (R_a_)** consists of overstorey leaf (R_ol_), stem (R_stem_), root (R_root_), understorey aboveground (R_ua_) (Figure S17), and growth respiration (R_grow_) (Figure S18). R_ol_ and R_ua_ were based on MAESPA simulation (Figure S17a, c), as described in the respective GPP sections. R_grow_ was estimated by taking a constant fraction of 30% of total NPP as measured directly on *E. tereticornis* trees^56^.

R_stem_ was estimated from measurements of stem CO_2_ efflux performed in three dominant trees per plot (Figure S17b). Collars were horizontally attached to the stem at an approximate height of 0.75 m, and R_stem_ was measured with a portable infrared gas analyzer coupled to a soil respiration chamber adapted for this purpose^57^. Measurement campaigns were performed every one or two months from December 2017 to October 2018, and the relationship between R_stem_ and air temperature (T_air_) was used to extrapolate R_stem_ across the surveyed period, following R_stem_ = 0.1866 * 2.84^Tair/10^ (r^2^ = 0.42, p < 0.0001). R_stem_ was then upscaled to the stand level considering the ratio of trunk stem axial surface per unit of soil surface measured per plot. Stem surface area was directly inferred from the Terrestrial Laser Scanning (TLS) data through quantitative structure models presented in Ref. 58 and 59. TLS data were acquired with a RIEGL VC-400 terrestrial laser scanner (RIEGL Laser Measurement Systems GmbH). Stem surface area was derived from the TLS data following a two-step approach: (i) manually extracting single tree from the registered TLS point cloud; and (ii) deriving parameters for an extracted single tree. Once a tree is extracted from the point cloud, the next step was to strip off the leaves, and segment the point cloud into stem and branches. Finally, the surface of the segments was reconstructed with geometric primitives (cylinders). The method used a cover set approach, where the point cloud was partitioned into small subsets, which correspond to small connected patches in the tree surface.

R_root_ was partitioned into fineroot (R_froot_) and coarse root (R_croot_) respiration (Figure S17d). Both R_froot_ and R_croot_ were estimated based on soil temperature at 20 cm depth. Mass-based rates of R_froot_ were obtained from measured rates in seedlings of *E. tereticornis*^60^. R_croot_ was estimated using a proxy based on measured rates of wood respiration of branches (c. 7 mm diameter) in trees (8 to 9 m height) of *E. tereticornis*^61^. The equations are:

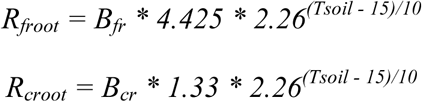

where *R_froot_* and *R_croot_* are fine root and coarse root respiration rates, respectively, *T_soi_*_l_ is soil temperature at 15 min interval, *B_fr_* and *B_cr_* are fineroot and coarse root biomass, respectively. Here we assumed fraction of coarse root at top 30 cm of soil is 60 % to represent coarse root respiration at this soil profile.

**Carbon efflux due to insect respiration (R_ins_)** was estimated as the net difference between NPP_ins_ and Frass, assuming no net change in insect biomass (Figure S16b).

**Soil respiration (R_soil_):** The rate of soil CO_2_ efflux was measured at eight locations within each plot, where a permanent PVC collar inserted into the soil was co-located with soil TDR probes for continuous measurements of soil temperature (5-cm-depth) and volumetric water content (0 to 21-cm-depth; CS650-L; Campbell Scientific, Logan, UT, USA). R_soil_ was measured manually at all collar locations every 2-3 weeks, in addition to 30-minute measurements using automated chambers (Li-8100-103; Licor) at one location within each plot, resulting in >300,000 observations over the study period^24^. These data were used to parameterize a semi-mechanistic model of R_soil_, in which R_soil_ was predicted based on measurements of soil properties, soil physics, and measured soil temperature and volumetric water content^62^. This model successfully recreated the observed fluxes (r^2^ between predicted and observed survey R_soil_ was 0.65)^24^. Annual sums of R_soil_ were derived by summing the averaged daily fluxes over eight locations within each plot, where daily fluxes at each location were predicted based on the semi-mechanistic model and daily soil temperature and volumetric water content data taken adjacent to each measurement collar. Soil heterotrophic respiration (R_hetero_) was taken as the net difference between R_soil_ and R_root_ (Figure S19). Total ecosystem respiration (R) was calculated as the sum of R_a_, R_hetero_, R_ins_, and VC.

**Volatile carbon (VC;** Figure S20) flux as isoprene (C_5_H_8_) was estimated using the Model of Emissions of Gases and Aerosols from Nature (MEGAN)^63^. Isoprene represents over half of all VOC species emitted by vegetation globally. A MEGAN box-model was built from the version used in Ref. 64, centered on the EucFACE facility to calculate hourly emissions of isoprene across the period 2013-2016 for all six plots:

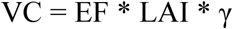

Where EF is the isoprene basal emission factor, γ is the emission activity factor, accounting for changes in the isoprene response due to light, temperature, leaf age and soil moisture. The MEGAN simulations were driven by daily input data of LAI, soil moisture, and hourly input data of photosynthetic active radiation, temperature, atmospheric pressure, wind speed and relative humidity. The isoprene EFs were measured as 6.708 mg m^-2^ h^-1^ for ambient CO_2_ plots and 5.704 mg m^-2^ h^-1^ for elevated plots. The EFs were derived from in-line photosynthetic gas-exchange measurements coupled with simultaneous volatile isoprenoid sampling. The isoprene emissions were collected in sterile stainless steel thermal desorption tubes at the same time as gas exchange was measured, and these were capped and later thermally desorbed for off-line volatile analysis in the laboratory using a Shimadzu GC/MS. The chromatographic peaks were identified by comparing them to isoprene standards and reference mass spectra in the NIST Mass Spectral Library (https://www.nist.gov/srd). The box-model produced isoprene was converted to carbon content using the molecular weight ratio of carbon to isoprene.

#### Net Ecosystem Production

Net ecosystem production (NEP) was estimated based on three different methods that estimated NEP in relatively independent ways (Figure 3), similar to Ref. 65. The first method considered NEP as the difference between total ecosystem influx and total ecosystem outflux (i.e. In - Out), which relied on both process-based modeling and empirical upscaling of respiratory fluxes collected from the field. The second method considered NEP as NPP minus R_hetero_ (i.e. NPP - R_hetero_), with NPP relying mostly on litter-based production estimates, and R_hetero_ relying on R_soil_ and R_root_ estimates. The third method considers NEP as the sum of changes in carbon pools in the ecosystem (i.e. ΔC_pools_), which was mostly determined by biomass estimates. Equations for each method are provided below:

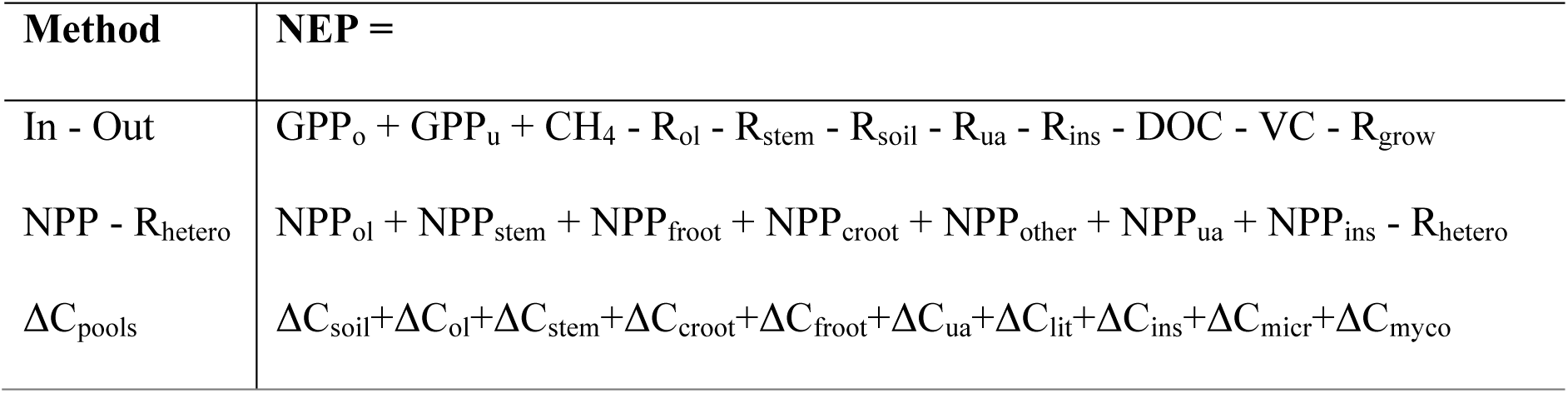

| Method | NEP = |
| --- | --- |
| In - Out | $\text{GPP}_o + \text{GPP}_u + \text{CH}_4 - R_{ol} - R_{stem} - R_{soil} - R_{ua} - R_{ins} - \text{DOC} - \text{VC} - R_{grow}$ |
| $\text{NPP} - R_{\text{hetero}}$ | $\text{NPP}_{ol} + \text{NPP}_{stem} + \text{NPP}_{froot} + \text{NPP}_{croot} + \text{NPP}_{other} + \text{NPP}_{ua} + \text{NPP}_{ins} - R_{\text{hetero}}$ |
| $\Delta C_{\text{pools}}$ | $\Delta C_{soil} + \Delta C_{ol} + \Delta C_{stem} + \Delta C_{croot} + \Delta C_{froot} + \Delta C_{ua} + \Delta C_{lit} + \Delta C_{ins} + \Delta C_{micr} + \Delta C_{myco}$ |

### Carbon budget evaluation

We evaluated the mass balance of our estimated ecosystem carbon budget in two ways. Firstly, we compared model simulated GPP with the aggregated sum of NPP and R_a_ (Extended Data Figure 2a, b). GPP was simulated by a stand-level ecophysiological model, driven by hourly meteorological data and parameterized with site-specific ecological data^19^. This GPP should equal to the aggregation of NPP (NPP_ol_ + NPP_stem_ + NPP_froot_ + NPP_croot_ + NPP_other_ + NPP_ua_ + NPP_ins_) and R_a_ fluxes (R_ol_ + R_stem_ + R_root_ + R_ua_ + R_grow_), which were mostly extrapolated based on field data. Secondly, R_soil_ estimated based on soil collar flux measurements^23^ was evaluated against the sum of litterfall and R_root_ (Extended Data Figure 2c, d), assuming minimal changes in soil carbon stock (as change over this short period of time is beyond the detection limit in a complex and slow-growing mature forest ecosystem like EucFACE). Here, litterfall was the sum of NPP_ol_ + NPP_froot_ + NPP_croot_ + NPP_other_ + NPP_ua_ + Frass, and R_root_ was extrapolated based on root biomass and temperature functions.

### Statistical analyses

We performed linear mixed-model analysis using the “lmer” function within the “lme4” package^66^ in software R^67^ to determine the CO_2_ treatment effect on all reported variables. All fluxes were reported at an annual rate (gCm^-2^yr^-1^). In our model, date and CO_2_ treatment were considered as fixed factors, plot as a random factor, and plot-specific pre-treatment LAI (i.e. 4-month average LAI before full CO_2_ treatment was switched on) as a covariate to account for pre-treatment differences among treatment plots. Normalizing all response variables with a covariate that integrates light, water and nutrient constraints helps to isolate the CO_2_ effect^21^, as has been done previously at the site^22^ and elsewhere^8,21^. Confidence intervals for the CO_2_ effect size of individual variables were reported using the function “confint”, which applies quantile functions for the t-distribution after model fitting. Confidence intervals for the predicted flux and pool were reported as the standard deviation of the plot-specific totals (n = 3). Similarly, confidence intervals for the aggregated fluxes (e.g. NPP) were reported by summing individual component fluxes that constituent the aggregated flux for each plot and computing the standard deviations across plots (n = 3). Finally, confidence intervals for the CO_2_ effect size (SD_agg_) of some aggregated fluxes (e.g. NPP) were calculated by pooling the standard deviations of the aggregated fluxes for ambient (SD_amb_) and elevated CO_2_ treatment (SD_ele_), following:

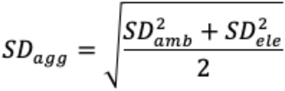

## Data statement

Data and code will be made available via Research Data Australia upon acceptance of the manuscript.

## Acknowledgements

EucFACE was built as an initiative of the Australian Government as part of the Nation-building Economic Stimulus Package, and is supported by the Australian Commonwealth in collaboration with Western Sydney University. We acknowledge the technical support by V. Kumar and C. McNamara, and the team of people who have assisted with data collection. The Eucalyptus tree vector in Figure 1 is from Heydon, L. *Eucalyptus spp*. Integration and Application Network, University of Maryland Center for Environmental Science (ian.umces.edu/imagelibrary/). This work was partially supported by the following grants from the Australian Research Council: DP130102501 (to JRP and ICA), DP110105102 and DP160102452 (to DSE). RLS received funding from Research Foundation Flanders and the European Union’s Horizon 2020 research and innovation programme under the Marie Skłodowska-Curie grant agreement no. 665501. RO-H. is financially supported by a Ramón y Cajal Fellowship from MICIU (RYC-2017-22032).

## Author contributions

MJ, BEM, RAD and JED designed the synthesis, compiled the data, and performed the analyses. MJ, BEM, RAD, JED, ICA, CVMB, MMB, LC-G, YC, LC, KYC, SLF, ANG, TEG, SH, SNJ, CAM, KM, BDM, LN, UNN, NJN, RO-H, VSP, EP, JP, JRP, SAP, PBR, AAR, MR, PR, RLS, BKS, BS, MGT, JKMW, AW-K, JY and DSE collected data and contributed to data analyses. JY and BEM performed the MAESPA model simulations, with contributions from MGDK and RAD. JED and AAR performed soil respiration gap-filling and modelling. KME performed isoprene emission model simulation. MJ and LC-G conceptualized Figure 1, and LC-G implemented the graphic design. MJ wrote the initial manuscript, with significant input from BEM, JED, BS, PBR, SZ, MGDK, MGT and DSE. All authors edited and approved the manuscript.

---

**Extended Data Table 1.**
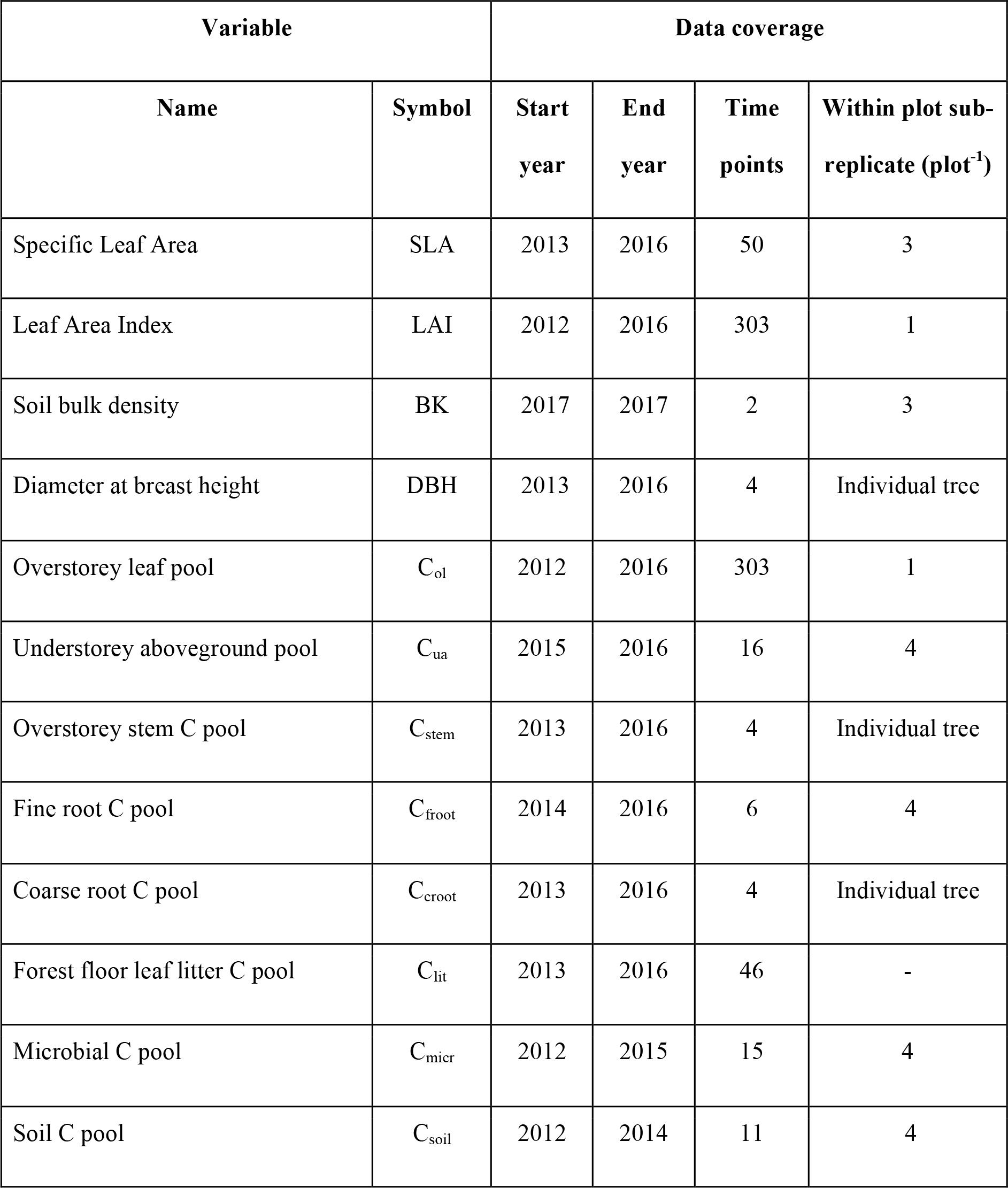
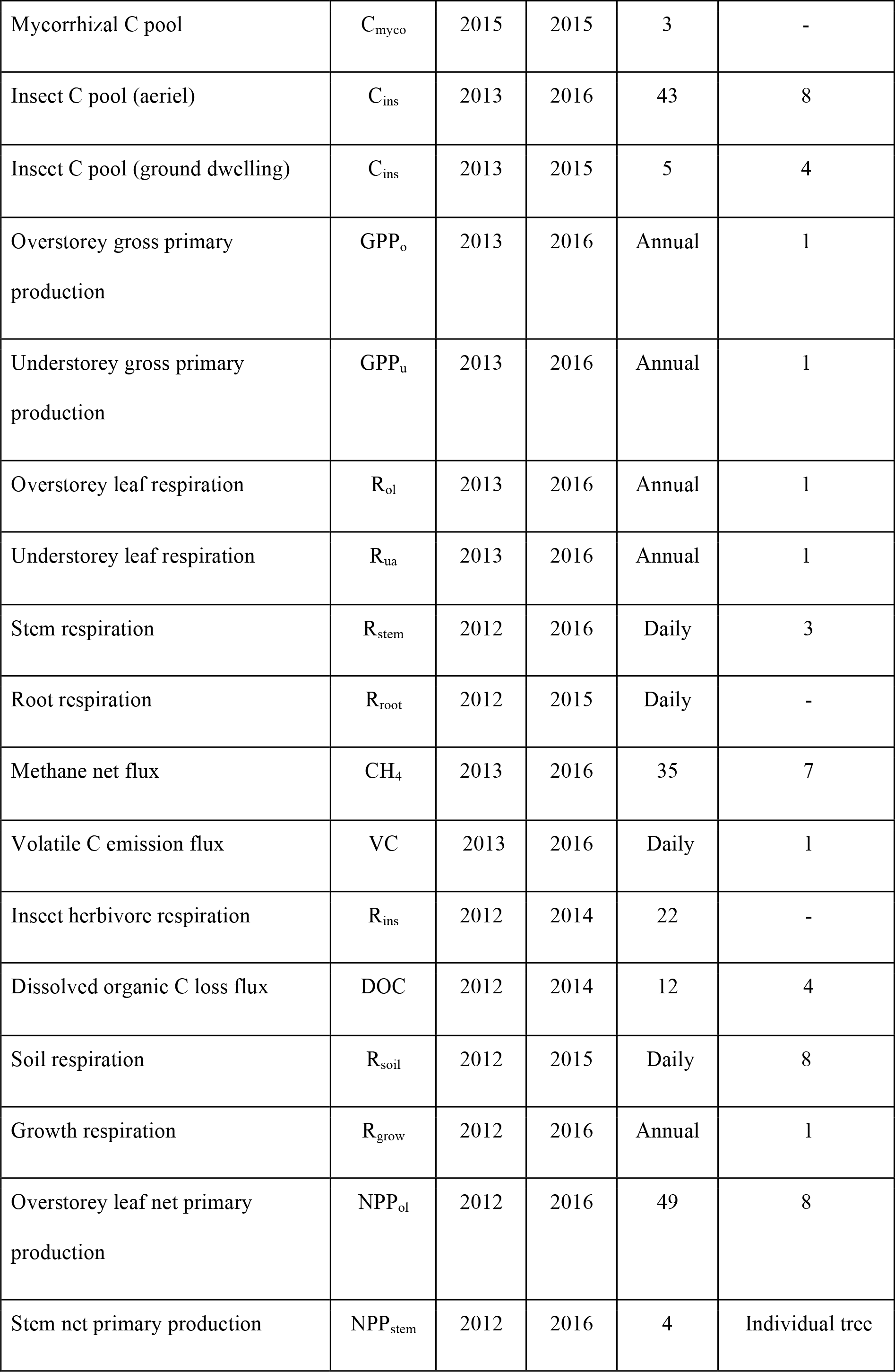
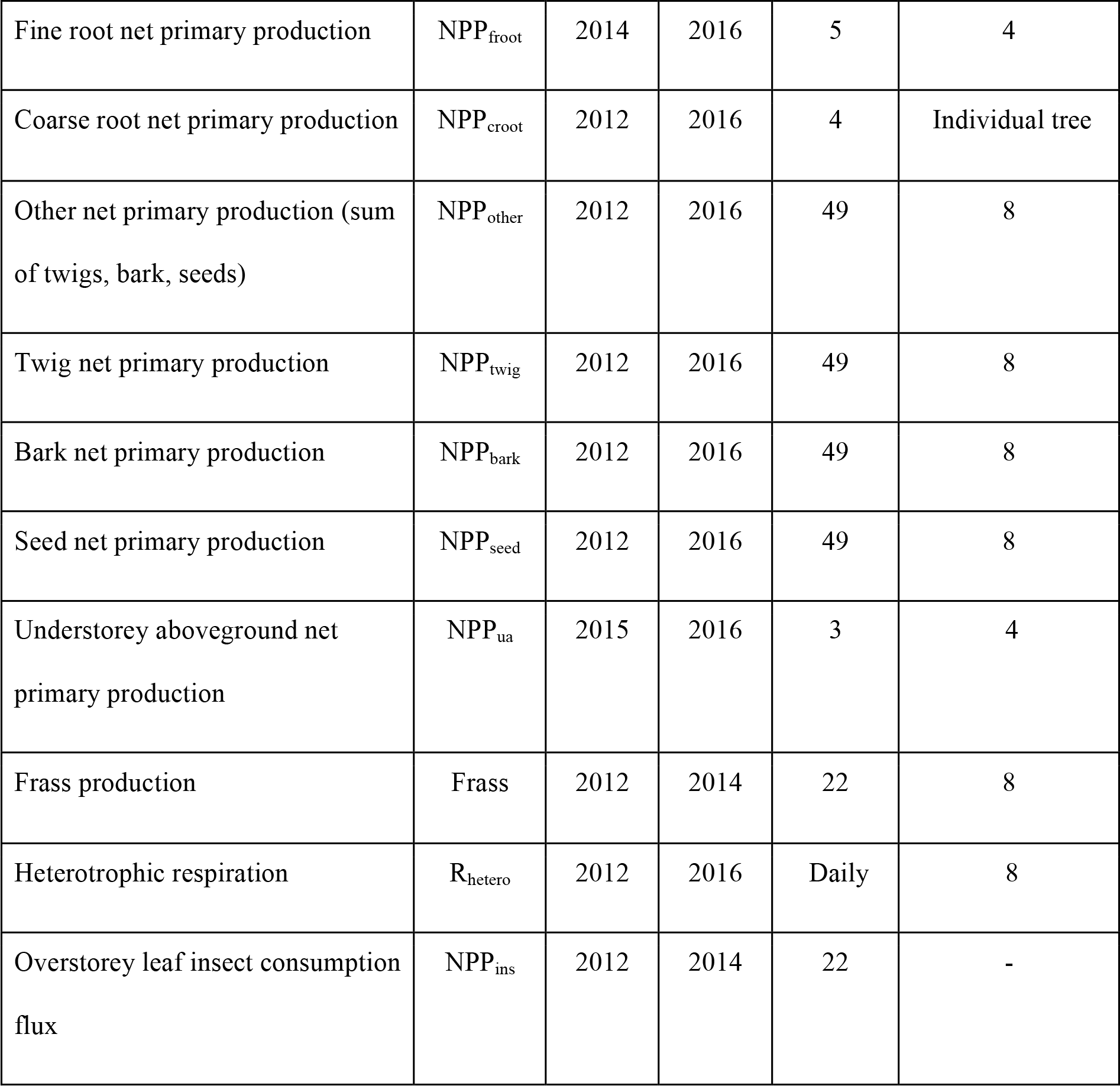
Definition and data availability of variables. Data availability includes start and end year of data included in this study. Time points indicate the number of data collections over the available data period. Within plot sub-replicate indicate the number of replicates within each treatment plot. The detailed methods for estimating each variable is provided in the Method section.

**Extended Data Table 2.** Carbon (C) fraction used to convert from biomass into C content.

| Variable | Symbol | Mean value | Data source |
| --- | --- | --- | --- |
| C fraction of overstorey leaf pool | $f_{ol}$ | 0.5 | EucFACE data |
| C fraction of understorey aboveground pool | $f_{ua}$ | 0.456 | EucFACE data |
| C fraction of stem pool | $f_{stem}$ | 0.445 (ambient plots)<br>0.448 (elevated plots) | EucFACE data |
| C fraction of coarse root pool | $f_{croot}$ | 0.445 (ambient plots)<br>0.448 (elevated plots) | Assumed the same as $f_{stem}$ |
| C fraction of fine root pool | $f_{froot}$ | 0.40 (ambient plots)<br>0.42 (elevated plots) | EucFACE data |
| C fraction of overstorey leaflitter pool | $f_{lit}$ | 0.5 | EucFACE data |
| C fraction of aboveground insect pool | $f_{ins}$ | 0.5 | Ref 48 |
| C fraction of frass production | $f_{frass}$ | 0.53 | EucFACE data |
| C fraction of microbial pool | $f_{micr}$ | 0.534 (ambient plots)<br>0.493 (elevated plots) | EucFACE data |
| C fraction of mycorrhizal pool | $f_{\text{myco}}$ | 0.534 (ambient plots)<br>0.493 (elevated plots) | Assumed the same as $f_{\text{micr}}$ |
| C fraction of soil pool | $f_{\text{soil}}$ | 0.016 (ambient plots)<br>0.017 (elevated plots) | EucFACE data |
| C fraction of twigs, barks and seeds production | $f_{\text{other}}$ | 0.5 | Assumed |

**Extended Data Figure 1.**
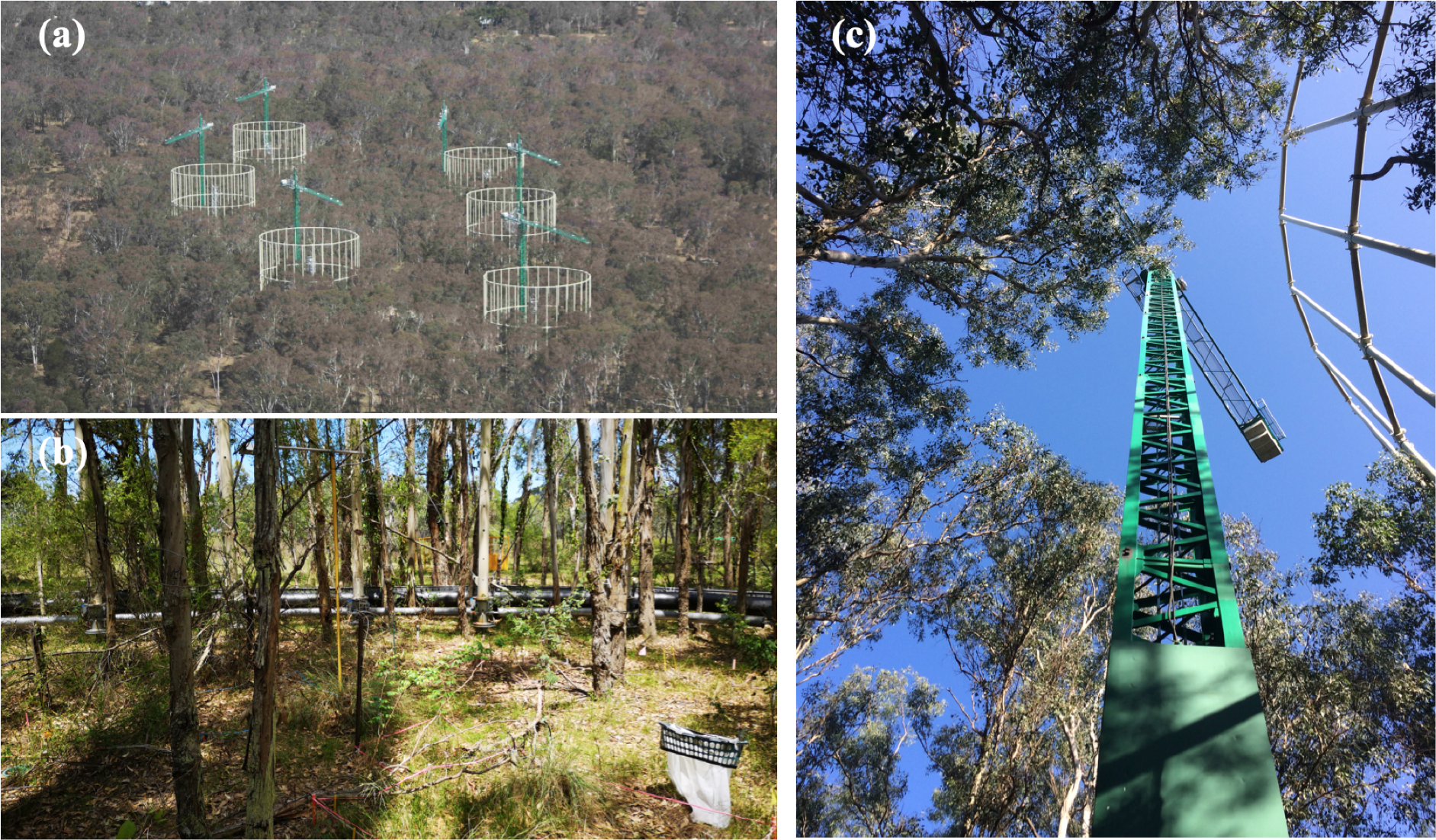
The *Eucalyptus* free air carbon dioxide enrichment experiment facility (EucFACE). **a)** A spatial overview of the forest and the facility (photo credit: David S. Ellsworth), **b)** an overview of the understorey vegetation and infrastructure inside a plot (photo credit: Mingkai Jiang), and **c)** a bottom-up look of the canopy structure and the crane (photo credit: Mingkai Jiang).

**Extended Data Figure 2.**
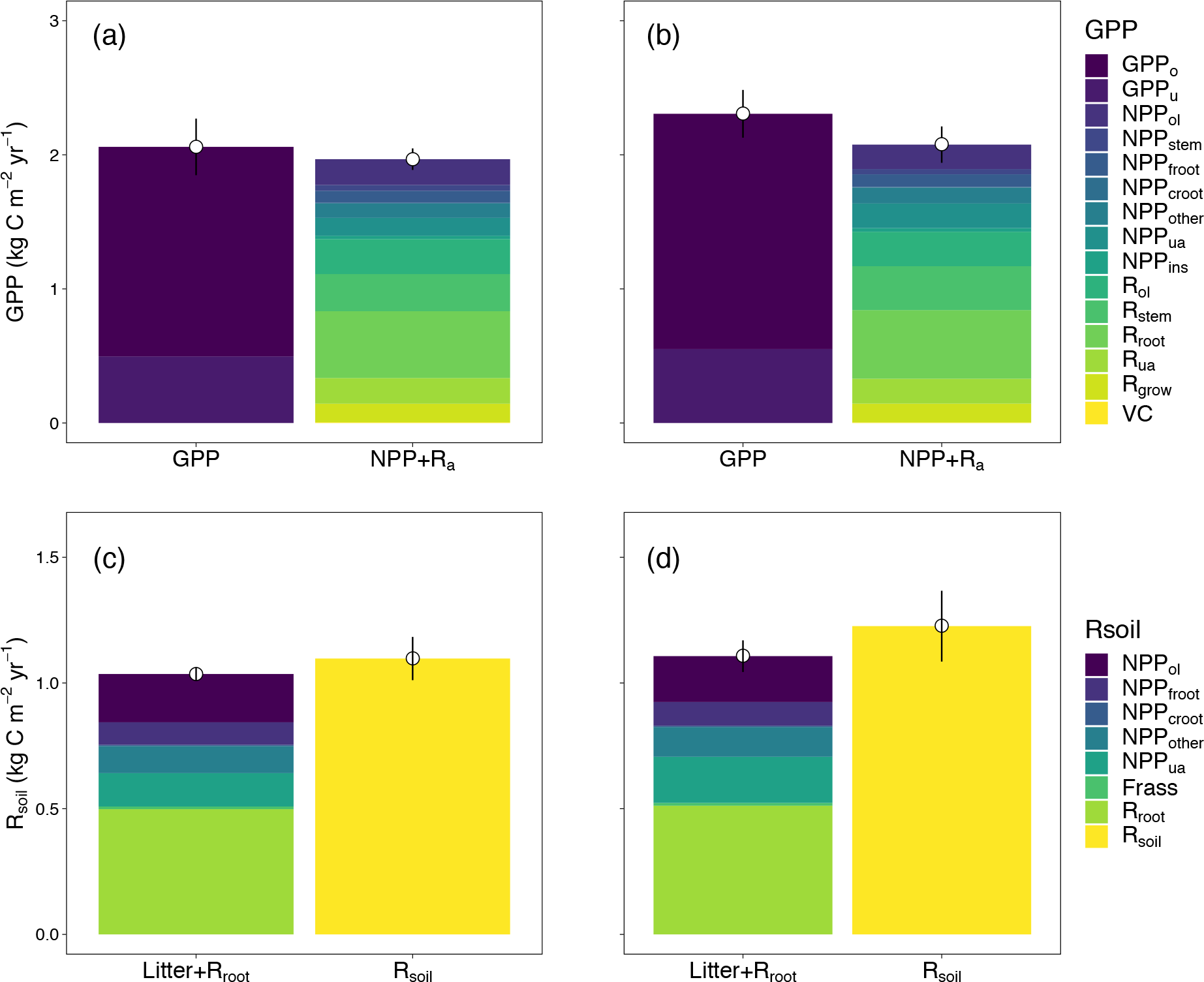
Estimates of (a and b) gross primary production (GPP) and (c and d) soil respiration (R_soil_) based on different methods for both (a and c) ambient and (b and d) elevated CO_2_ treatment at EucFACE. For estimates of GPP, we compared the model simulated total GPP of overstorey and understorey (GPP_o_ and GPP_u_, respectively), with the sum of data-driven estimates of net primary production (NPP) and autotrophic respiration (R_a_), which include NPP of overstorey leaf (NPP_ol_), stem (NPP_stem_), fineroot (NPP_froot_), coarse root (NPP_croot_), twigs, barks and seeds (NPP_other_), understorey aboveground (NPP_ua_), leaf consumption by insects (NPP_ins_), and respiratory fluxes of overstorey leaf (R_ol_), stem (R_stem_), root (R_root_), understorey aboveground (R_ua_), growth (R_grow_), and volatile carbon emission (VC). For estimates of R_soil_, we compared direct estimates of R_soil_ scaled up from soil chamber measurements, with the sum of litterfall and independent estimates of root respiration (Litter + R_root_), assuming no net change in soil carbon stock over time. Here litterfall was inferred based on NPP of overstorey leaf (NPP_ol_), fineroot (NPP_froot_), coarse root (NPP_croot_), twigs, barks and seeds (NPP_other_), understorey aboveground (NPP_ua_), and frass production (Frass). These evaluations provide independent mass balance checks of the estimated ecosystem carbon budget. Each color represents a flux variable. Dotted point and vertical line represent treatment mean and standard deviation based on plot-level estimates of the aggregated flux (n=3). Values were normalized by a linear mixed-model with pre-treatment leaf area index as a covariate to account for pre-existing differences.

**Extended Data Figure 3.**
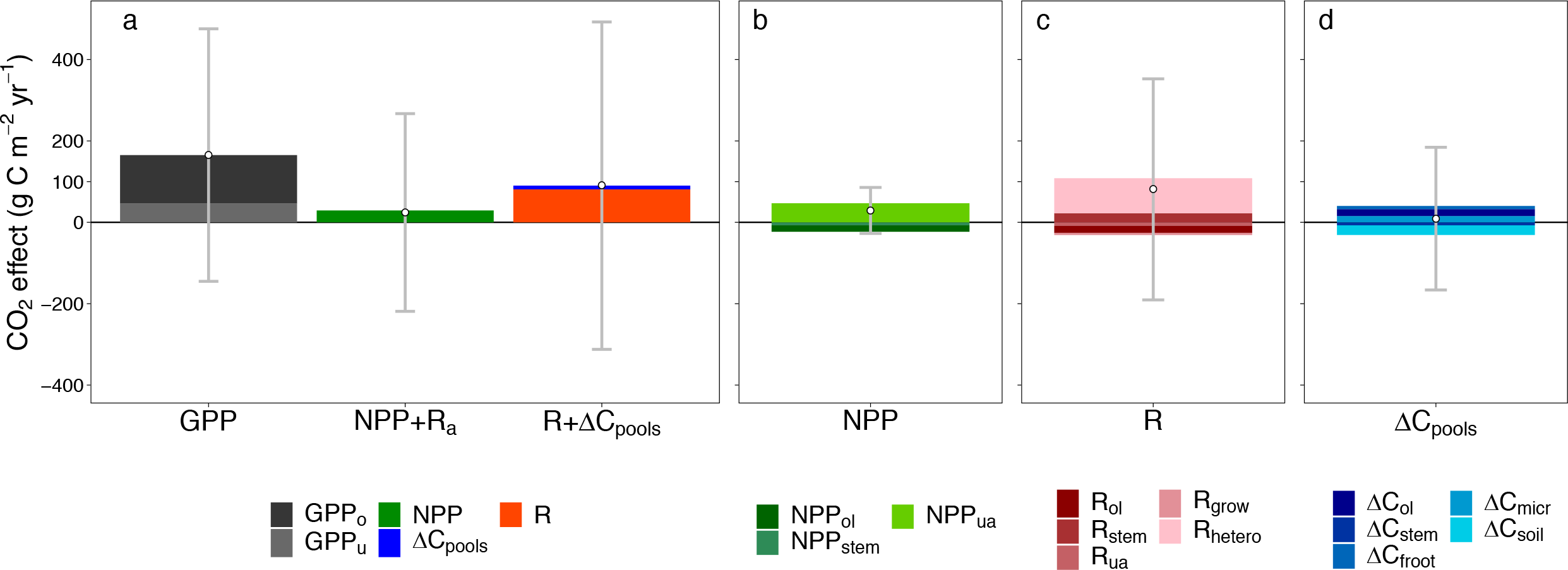
The fate of additional carbon fixed under elevated CO_2_ (eCO_2_) in a mature forest ecosystem (non-normalized analysis case). **a)** Column **“**GPP” represents the total eCO_2_ induced increase in overstorey and understorey gross primary production (GPP_o_ and GPP_u_, respectively), column “NPP + R_a_” represents the sum of net primary production and autotrophic respiration eCO_2_ response, and column “R + ΔC_pools_” represents the sum of ecosystem respiration and carbon storage eCO_2_ response. **b)** The relative contributions of individual NPP fluxes to the aggregated NPP response to eCO_2_, including overstorey leaf (NPP_ol_), stem (NPP_stem_), and understorey aboveground (NPP_ua_). **c)** The relative contributions of individual respiratory fluxes to the aggregated R response to eCO_2_, including overstorey leaf (R_ol_), stem (R_stem_), understorey aboveground (R_ua_), growth (R_grow_), and heterotroph (R_hetero_). **d)** The relative contributions of individual change in carbon storage to the aggregated ΔC_pools_ response to eCO_2_, including overstorey leaf (ΔC_ol_), stem (ΔC_stem_), fineroot (ΔC_froot_), microbe (ΔC_micr_), and soil (ΔC_soil_). Variables with an average CO_2_ effect of < 5 gCm^-2^yr^-1^ were excluded from the figure for better visual clarification. Each color represents a flux variable, point indicates the net sum of all variables for a column, and the grey confidence interval represents plot-level standard deviation (n=3) of the estimated column sum.

**Extended Data Figure 4.**
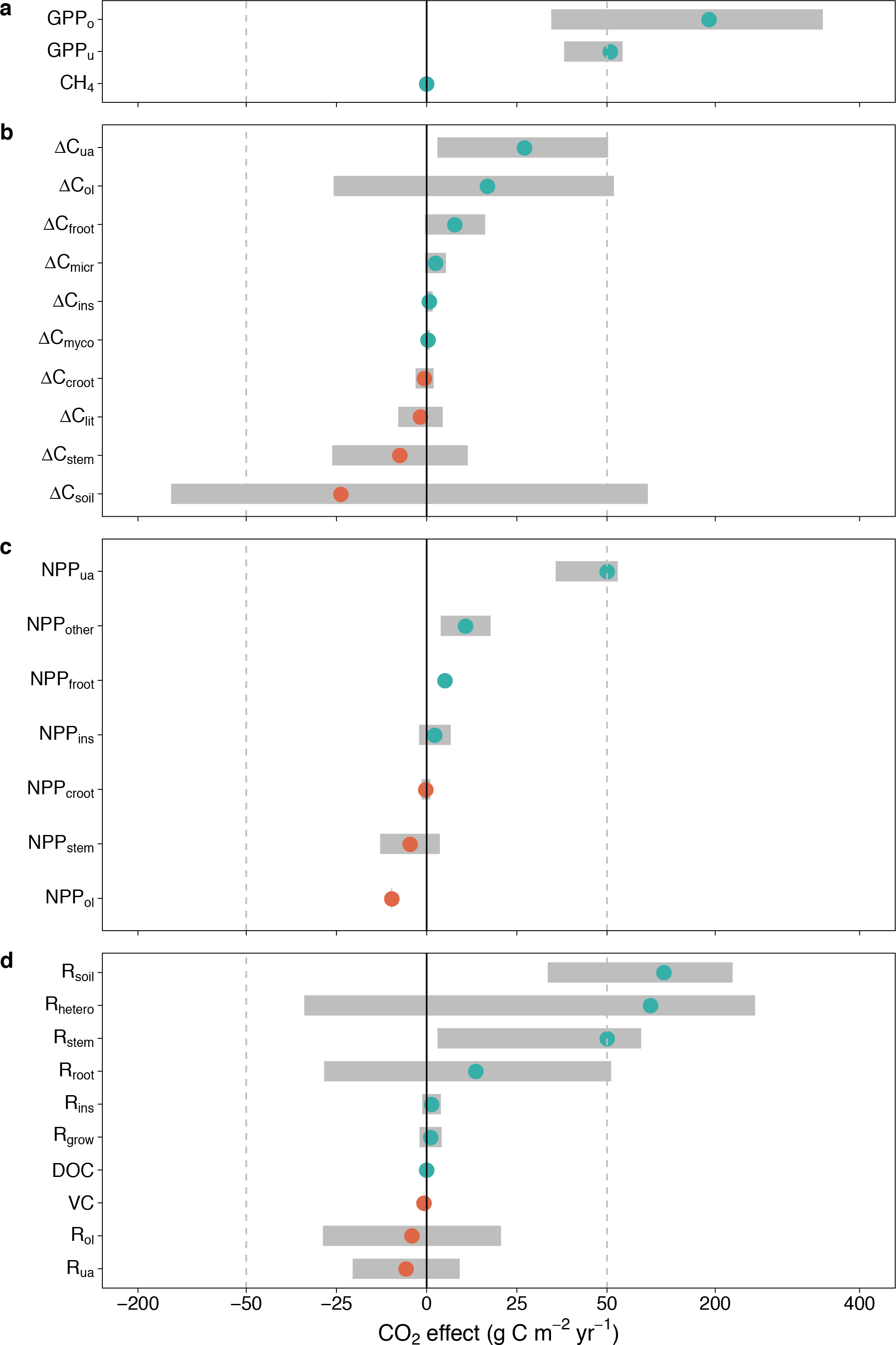
CO_2_ treatment effect (gCm^-2^yr^-1^) for all ecosystem fluxes at EucFACE. **a)** The CO_2_ response of gross ecosystem carbon uptake, including gross primary production of overstorey (GPP_o_) and understorey (GPP_u_), and soil methane uptake (CH_4_). **b)** The eCO_2_ response of annual incremental change in carbon pool (ΔC_pools_), including overstorey leaf (ΔC_ol_), stem (ΔC_stem_), coarse root (ΔC_croot_), fineroot (ΔC_froot_), understorey aboveground (ΔC_ua_), leaf litter (ΔC_lit_), soil (ΔC_soil_), microbe (ΔC_micr_), aboveground insect (ΔC_ins_), and mycorrhizae (ΔC_myco_). **c)** The eCO_2_ response of net primary production (NPP), including overstorey leaf (NPP_ol_), stem (NPP_stem_), coarse root (NPP_croot_), fineroot (NPP_froot_), understorey aboveground (NPP_ua_), twigs, barks and seeds (NPP_other_), and leaf insect consumption (NPP_ins_). **d)** The eCO_2_ response of ecosystem respiration (R) and other out-going flux, including respiration fluxes of overstorey leaf (R_ol_), stem (R_stem_), root (R_root_), understorey aboveground (R_ua_), growth (R_grow_), insect (R_ins_), heterotroph (R_hetero_), and soil (R_soil_), and volatile carbon emission (VC) and dissolved organic carbon leaching (DOC). Dots and grey bars represent means and standard deviations of the CO_2_ treatment difference, predicted by a linear mixed-model with plot-specific pre-treatment leaf area index as a covariate. Orange dots indicate negative means and light green dots indicate positive means. Dashed lines indicate change of scale along the x-axis.

